# A versatile CRISPR toolbox to study neurological Ca_V_3.2 channelopathies by promoter-mediated transcription control

**DOI:** 10.1101/2021.03.20.436250

**Authors:** Despina Tsortouktzidis, Anna R. Tröscher, Herbert Schulz, Susanne Schoch, Albert J. Becker, Karen M.J. van Loo

## Abstract

Precise genome editing in combination with viral delivery systems provides a valuable tool for neuroscience research. Traditionally, the role of genes in neuronal circuits has been addressed by overexpression or knock-out/knock-down systems. However, those techniques do not manipulate the endogenous loci and therefore have limitations. Those constraints include that many genes exhibit extensive alternative splicing, which can be regulated by neuronal activity. This complexity cannot be easily reproduced by overexpression of one protein variant. The CRISPR activation and interference/inhibition systems (CRISPRa/i) directed to promoter sequences can modulate the expression of selected target genes in a highly specific manner. This strategy could be particularly useful for the overexpression of large proteins and for alternatively spliced genes, e.g. for studying large ion channels known to be affected in ion channelopathies in a variety of neurological diseases. Here, we demonstrate the feasibility of a newly developed CRISPRa/i toolbox to manipulate the promoter activity of the *Cacna1h* gene. Impaired, function of the low-voltage-activated T-Type calcium channel Ca_V_3.2 is involved in genetic/mutational as well as acquired/transcriptional channelopathies that emerge with epileptic seizures. We show CRISPR-induced activation and inhibition of the *Cacna1h* locus in NS20Y cells and primary cortical neurons, as well as activation in mouse organotypic slice cultures. In future applications, the system offers the intriguing perspective to study functional effects of gain-of-function or loss-of-function variations in the *Cacna1h* gene in more detail. A better understanding of Ca_V_3.2 channelopathies might result in a major advancement in the pharmacotherapy of Ca_V_3.2 channelopathy diseases.

## 1 Introduction

The human genome encodes approximately 400 ion channel genes, encompassing both voltage-gated and ligand-gated ion channels (Hutchings et al., 2019). Ion channels are pore forming membrane proteins that allow ionic flows across membranes and are crucial for normal functioning of many tissues, including the central and peripheral nervous system, heart, kidney and liver (Cannon, 2007). Ion channel dysfunctions, also coined as ion ‘channelopathies’, have been associated with a large variety of diseases. Besides disorders of the nervous system (e.g. epilepsy, ataxia, Alzheimer’s disease and Autism spectrum disorders), also cardiac arrhythmia and several muscle, endocrine and renal disorders are linked to dysfunction of ion channels (Wang et al., 1996; Heeringa et al., 2009; Ryan et al., 2010). Still, many molecular and structural mechanisms of how channelopathies convert cells from a health to disease state are not fully understood, including critical time-windows during development as well as the potential reversibility by reconstituting normal expression / function of affected molecules. Major obstacles to manipulate ion channels are given by their large size, which limits options for widespread and *in vivo* overexpression and their diversification by alternative splicing.

Genetic editing using a modified CRISPR (clustered regularly interspaced palindromic repeats) system could be a powerful approach for the manipulation of large proteins that cannot be done by conventional techniques. Recently, substantial progress has been made in genomic editing by using specific applications of the CRISPR technique, including the CRISPR *activation* (CRISPRa) and CRISPR *interference/inhibition* (CRISPRi) technology. CRISPRa uses a catalytically dead Cas9 (dCas9) enzyme fused with a highly efficient transcriptional activator complex consisting of the tripartite transcriptional activator VP64-p65-Rta (VPR), shown to increase target gene expression even up to 320-fold (Chavez et al., 2015). CRISPRi also uses the dCas9 enzyme and is fused with the transcriptional repressor KRAB (Krüppel-associated box) protein, which results in up to 60-80% reduction in the expression of endogenous eukaryotic genes (Gilbert et al., 2013). Using these enhanced CRISPR technology systems, the expression of genes can be modified in their native context with utmost precision. Therefore, this strategy will be particularly useful for (a) the overexpression of large proteins, which is difficult to accomplish by conventional techniques and (b) for alternatively spliced genes.

In this study, we have developed a CRISPRa/i toolbox for manipulating the expression of a well-described ion channelopathy gene, *Cacna1h*, encoding the low-voltage-activated T-Type calcium channel Ca_V_3.2. Ion channelopathies for *Cacna1h* have been described particularly for epilepsy variants (Khosravani et al., 2004, 2005; Powell et al., 2009; Souza et al., 2019), but were also reported for other neurological diseases including autism spectrum disorders (Splawski et al., 2006), amyotrophy lateral sclerosis (ALS; (Rzhepetskyy et al., 2016)) and pain disorders (Souza et al., 2016). By using the unique CRISPRa/i toolbox, we demonstrate to specifically modulate *Cacna1h* gene expression in different cell types in order to closely recapitulate Ca_V_3.2 channelopathies.

## 2 Material and Methods

### 2.1 Design of sgRNAs

sgRNA design was performed using the computational software Benchling (Cloud-Based Informatics Platform for Life Sciences R&D | Benchling, 2021). The previously validated *Cacna1h* promoter (Van Loo et al., 2012) was used to set the sgRNAs targeting sequence. The sgRNAs were selected based on their local off-target scores, proximity to the start-ATG, a distance of at least 50 bp between each other and a 100% match in targeting both the mouse and rat genome.

### 2.2 Cloning

In order to generate the pAAV-U6-sgRNA plasmids, we first cloned a U6-BbsI/BbsI cassette into the pAAV-Syn-MCS backbone. For this, the U6-BbsI/BbsI cassette was PCR amplified from px458 (Addgene #48138, (Ran et al., 2013)) (forward: 5’-ctgcggccgcacgcggagggcctatttcccatgattcc-3 and reverse: 5’-aattcaatcgatgcgggtacctctagagccatttgt-3’) and inserted by in-fusion cloning (Takara Bio Europe/Clontech) into pAAV-Syn-MCS digested with MluI/AsiSI. Next, the polyA was removed by digestion with AfeI/PmlI and subsequent self-ligation. The sgRNAs were annealed and cloned into the BbsI sites (Cloud-Based Informatics Platform for Life Sciences R&D | Benchling, 2021). Briefly, the annealing was performed using 10 µM of each oligo, 1X T4 ligation buffer and 5U of T4 PNK in a total volume of 10 µL, following an incubation at 37 °C for 30 minutes and 95 °C for 5 minutes, the reaction was cooled at room temperature. The annealed oligos (1 µL) were cloned using 25 ng of construct, 1X T4 ligation buffer, T4 ligase (200 U) and BbsI (2.5 U) in a total volume of 10 µL, the reaction was performed by 30 cycles at 37 °C for 5 minutes and 23 °C for 5 minutes. The primers for cloning the sgRNAs were as follow: sgRNA1-*Cacna1h*: forward 5’-caccggggcgtcgttcctgggcca-3’ and reverse 5’-aaactggcccaggaacgacgcccc-3’; sgRNA2-*Cacna1h*: forward 5’-caccgagagacaaagacatcccgg-3’ and reverse 5’-aaacccgggatgtctttgtctctc-3’; sgRNA-*Lac*Z: forward 5’-caccgtgcgaatacgcccacgcgat-3’ and reverse 5’-aaacatcgcgtgggcgtattcgcac-3’.

The all-in-one CRISPRa lentiviral system was generated by replacing the promoter of pLenti-Ef1a-dCas9-VPR (Addgene#114195, (Savell et al., 2019)) by the human synapsin promoter using in-fusion cloning. For this, pAAV-Syn-MCS was used as PCR template for the synapsin promoter (forward 5’-gtttggttaattaagagtgcaagtgggttttaggacc-3’, and reverse 5’-atagtcggtggcagcatcgatgcgatcgcatgc-3’) and pLenti-Ef1a-dCas9-VPR was digested with AfeI/KpnI. In a second step, the *Cacna1h* and *Lac*Z sgRNA sequences from the pAAV-U6-sgRNA plasmids (see above) were cloned by in-fusion cloning into the PacI site of pLenti-syn-dCas9-VPR (forward: 5’-gatccagtttggttagagggcctatttcccatgattcct-3’ and reverse: 5’-ccacttgcactcttactagagccatttgtctgcagaat-3’), resulting in pLenti-U6-sgRNA_(Cacna1h/LacZ)_-Syn-dCas9-VPR.

The all-in-one CRISPRi lentiviral system was produced by replacing the promoter and U6 cassette of pLV-hU6-sgRNA-hUbC-dCas9-KRAB-T2a-eGFP (Addgene#71237, (Thakore et al., 2015) for the synapsin promoter by in-fusion cloning. The backbone was digested with XbaI/Pac and the synapsin promoter PCR amplified from pAAV-Syn-MCS (forward: 5’-gatccagtttggttaattaagtagactgcagagggccctg-3’, reverse: 5’-ccatggtggctctagagaattcaatcgatgcgatcgcatgcgc-3), resulting in pLenti-syn-dCas9-KRAB-T2A-eGFP. Subsequently, the U6-sgRNA_Cacna1h/LacZ_ cassettes from the pAAV-U6-sgRNA plasmids (see above) were inserted into the PacI site of pLenti-syn-dCas9-KRAB-T2A-eGFP by in-fusion cloning (forward: 5’-gatccagtttggttagagggcctatttcccatgattcct-3’, reverse: 5’-tctgcagtctacttagcgcacgcgctaaaaacggact-3’).

### 2.3 Cell culture, transfection and luciferase assay

NS20Y cells (Sigma, 08062517) were maintained at 37 °C and 5 % CO2 in DMEM (Sigma, D6546) supplemented with 10 % (v/v) heat inactivated FBS, 2mM L-Glutamine, 100 units/mL penicillin/streptomycin. Cells were transfected in 24 well plates using Lipofectamine (Invitrogen) following the manufacturer’s instructions. The DNA concentration used per well: *Cacna1h*-Luciferase or *Cacna1h*-mRuby 100 ng, CMV-VPR (Addgene# 63798, (Chavez et al., 2015)) or CMV-KRAB (Addgene#110821, (Yeo et al., 2018)) 200 ng, pAAV-sgRNA 200 ng and hypB-CAG-2A-eGFP 100 ng. Following 48 hours after transfection, the cells were imaged or collected for luciferase assays. Luciferase assays were performed using the Dual Luciferase Reporter Assay System (Promega) according to the manufacturer’s specifications. Firefly luciferase activity was determined using the Glomax Luminometer (Promega).

### 2.4 Viral production and neuronal transduction

AAV1/2 viruses were produced by large-scale triple CaPO4 transfection of HEK293-AAV cells (#240073, Stratagene) as described previously (Van Loo et al., 2012). Lentiviruses were produced in HEK293T cells using a second-generation lentiviral packaging system. The procedure was performed as described before (Van Loo et al., 2019). Briefly, 3 × 10^6^ HEK293T cells were transfected using GenJet transfection reagent (Signagen) with 7.5 µg psPax2 (Addgene #12260), 5 µg VSV-G (pMD2.G, Addgene #12259) and 3.3 µg of plasmid of interest. The medium was replaced 12 h after transfection with DMEM containing Glutamax (Invitrogen) and supplemented with 10 % FBS. After 72 h, the supernatant containing the viral particles was filtered using 0.45 PVDF membrane filters (GE Healthcare). The viral purification was performed using OptiPrep solution (Sigma-Aldrich) and centrifugation at 24,000 rpm for 2 h at 4°C. The isolated viral particles were mixed with TBS-5 buffer (50 mM Tris-HCl, 130 mM NaCl, 10 mM KCl, and 5 mM MgCl_2_). Viral particles were centrifuged at 24000 rpm, 2 h, 4 °C and resuspended in 15 µL TBS-5 Buffer. Dissociated primary neurons were prepared from mouse cortex (C57Bl6/N) at embryonic day 15 to 19 as described before (Woitecki et al., 2016). All animals were handled according to government regulations as approved by local authorities (LANUV Recklinghausen). All procedures were planned and performed in accordance with the guidelines of the University Hospital Bonn Animal-Care-Committee as well as the guidelines approved by the European Directive (2010/63/EU) on the protection of animals used for experimental purposes. Cells were kept in BME medium (Gibco) supplemented with 1% FBS, 0.5mM L-glutamine, 0.5% glucose and 1X B27 at 37°C and 5% CO2. Neuronal transduction (4 µL of virus/well) was performed at days in vitro (DIV) 4 in 24 well plates containing 70.000 cells/well. Cells were collected on DIV15 in lysis/binding buffer (Invitrogen, A33562) and stored at -80 °C until mRNA extraction.

### 2.5 RNA isolation and real time RT-PCR

RNA isolation and cDNA production was performed using Dynabeads mRNA Direct Micro Kit (Invitrogen, 61021) and RevertAid H Minus First strand cDNA Synthesis Kit (Thermo Fisher Scientific, K1632) following the manufacturer’s instructions. Quantitative PCR was performed in a Thermal Cycler (BioRad C1000 Touch, CFX384 Real-Time system). The reaction was performed using 1X Maxima SYBR Green/Rox qPCR Master Mix (Thermo Fisher Scientific, K0223), 0.3 µM of each primer and 1/10 synthesized cDNA (for NS20Y and 1/2 for Neurons) for a total volume of 6.25 µL. The qPCR conditions were as follow: 2 min at 50°C, 10 min at 95°C, 40 cycles of 15s at 95°C and 1min at 59°C. mRNA quantification was performed by real-time RT-PCR using the ΔΔCt-method. Quantification was based on synaptophysin (Chen et al., 2001). The primers used were Synaptophysin: forward 5-ttcaggactcaacacctcggt-3 and reverse 5-cacgaaccataggttgccaac-3; *Cacna1h*: forward 5-atgtcatcaccatgtccatgga-3 and reverse 5-acgtagttgcagtacttaagggcc-3; *Cacna1g*: forward 5-accctggcaagcttctctga-3 and reverse 5-gcggaggatgtacaccaggta-3; *Cacna1i*: forward 5-tcatccgtatcatgcgtgttct-3 and reverse 5-gggcccgcattcctgt-3; *Cacna1e*: forward 5-ggagtggatacccttcaatgagtc-3 and reverse 5-tctgttaccaccagagattgttgttc-3.

### 2.6 Mouse organotypic slices

Organotypic hippocampal slices were prepared from mice (C57Bl6/N) at postnatal day 3-6 as described before (Biermann et al., 2014). In brief, the hippocampus was isolated and cut in 350 µM thickness using a McIllwain tissue chopper. The slices were cultured in 6 well plates containing cell culture inserts 0.4 µM, 30 mm (Millipore) and medium (50% Neurobasal medium, 25% Hank’s balanced salt solution (without MgCl_2_, without CaCl_2_), 25% horse serum, 0.65% D(+)-glucose, 0.01 M Hepes, 2 mM L-glutamine and 0.5X B27). Cultured slices were kept at 37 °C, 5% CO_2_.

### 2.7 Cell imaging

Images were obtained using an inverted phase contrast fluorescent microscope (Zeiss Axio Observer A1 with objectives 20X, LD A-Plan and 5X, Fluar) and processed using Fiji. The cell surface area, set by the GFP fluorescence, was determined using the Weka trainable segmentation plugging. The integrated density, determined by the mRuby fluorescence was normalized to the cell surface area to obtain the reported values of integrated density/cell surface (IntDE/cell surfase).

### 2.8 Statistical analyses

Statistical analyses were performed using GaphPad software. One sample *t*-test, Student *t*-test and two way ANOVA followed by multiple comparison tests were used to compare significance of the results. Values were considered significant a *p* < 0.05. For all graphs data are displayed as mean ± SEM.

## 3 Results

To modulate the promoter activity of the *Cacna1h* gene, we first designed sgRNAs sequences targeting the mouse and rat *Cacna1h* gene. Using the Benchling computational tool, we selected two sgRNAs binding 66 (sgRNA1) and 131 (sgRNA2) base pairs (bp) upstream of the start-ATG of the *Cacna1h* gene (**Fig 1A**). Next, we tested the efficiency of the two sgRNAs to target the *Cacna1h* promoter and regulate its activity in neuroblastoma NS20Y cells. For this, we transfected the two sgRNAs together with (i) a minimal reporter unit expressing mRuby under control of the rat *Cacna1h* promoter, (ii) a ubiquitous promoter expressing eGFP and (iii) either a CRISPRa construct (dCas9-VPR, (Chavez et al., 2015)) or a CRISPRi construct (dCas9-KRAB, (Yeo et al., 2018)) in NS20Y cells and analyzed the fluorescence intensity 2 days after transfection (**Fig 1B, C**). A strong activation of the *Cacna1h* promoter was observed after co-transfection with the CRISPRa construct and an inhibition after co-transfection with the CRISPRi construct (**Fig 1C**). No modulation was observed for the ubiquitous CAG-eGFP construct, indicating that the sgRNAs only affected the activity of the *Cacna1h* promoter (**Fig 1C**).

**Figure 1.**
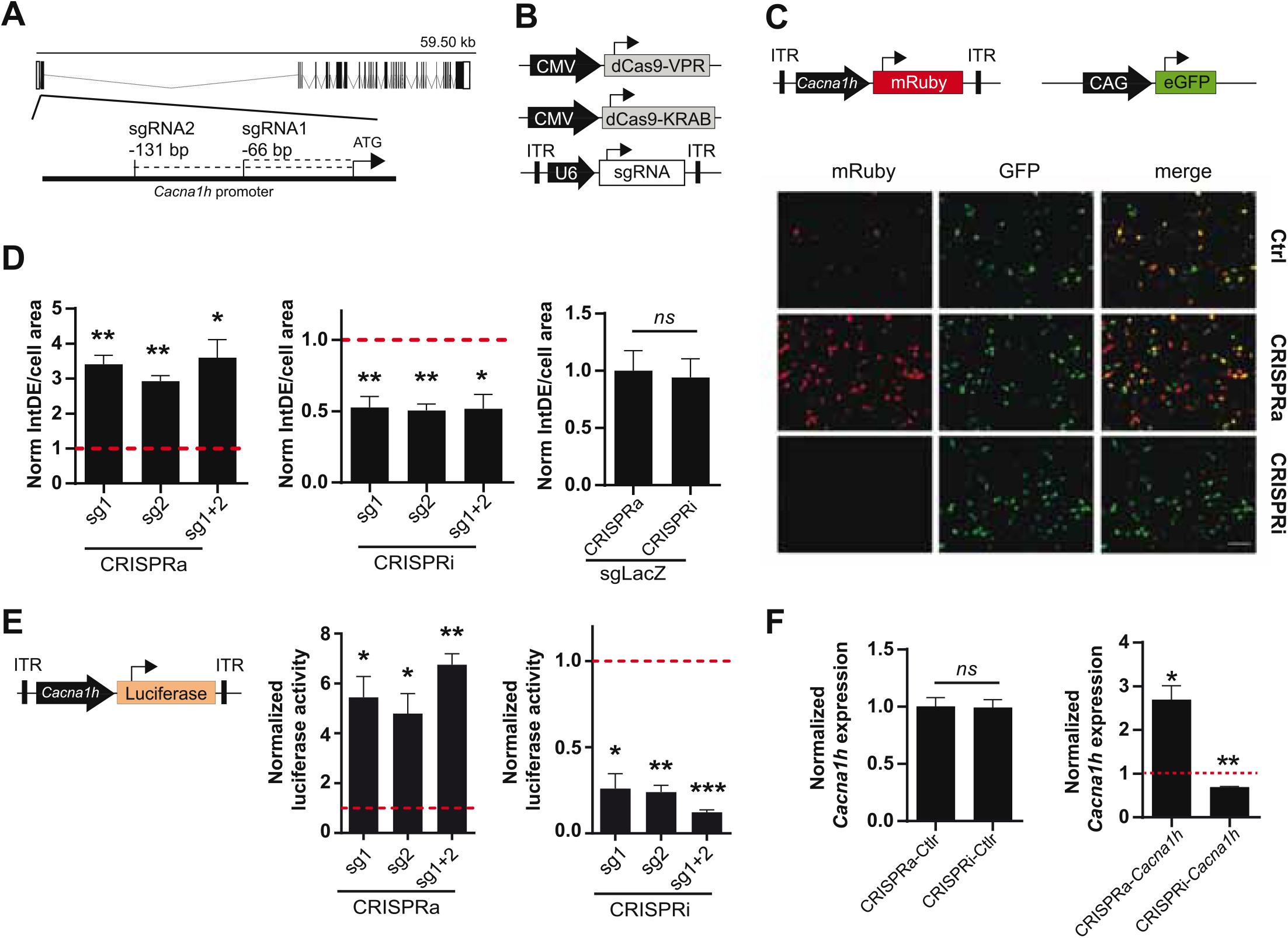
Validation of the CRISPRa/i system for modulating *Cacna1h* expression. **(A)** Schematic representation of the *Cacna1h* mouse gene. Exons and introns are shown as boxes and dash lines, respectively. The position of sgRNA1 and sgRNA2 are indicated respective to the start-ATG (bp: base pairs). **(B)** Schematic illustration of the CRISPRa (CMV-dCas9-VPR), CRISPRi (CMV-dCas9-KRAB) and the sgRNA (pAAV-U6-sgRNA) constructs. **(C)** NS20Y co-transfected with CRISPRa or CRISPRi, the two sgRNAs targeting *Cacna1h* (or *LacZ* as control), the reporter pAAV-*Cacna1h*-mRuby, and a CAG-eGFP construct as internal control for transfection. Scale bar = 100 µm. **(D)** Quantification of integrated density (IntDE) of mRuby per cell surface area (defined by CAG-eGFP positive cells). Values of IntDE/cell area were normalized to the controls of each treatment: CRSIPRa-*LacZ* for CRISPRa and CRISPRi-*LacZ* for CRISPRi (One sample t-test, N=4). The CRISPRa and CRISPRi controls (targeting *LacZ*) did not differ (data normalized to CRISPRa-*lacZ*, t-test, N=4). **(E)** Luciferase activity of cells co-transfected with pAAV-Cacna1h-luciferase, CRISPRa or CRISPRi and sgRNAs targeting *Cacna1h* (or *LacZ* as control). Data normalized to each control: CRISPRa-*LacZ* for CRISPRa treatment and CRISPRi-*LacZ* for CRISPRi treatment (One sample t-test, N=3). **(F)** *Cacna1h* mRNA expression in NS20Y cells co-transfected with CRISPRa or CRISPRi. Left panel: Cells transfected with CRISPRa and CRISPRi without sgRNAs did not show altered *Cacna1h* expression (N=3, t-test). Right panel: *Cacna1h* mRNA expression levels in NS20Y cells transfected with CRISPRa or CRISPRi and sgRNAs targeting *Cacna1h* (sgRNA1+2). Values were normalized to the mean of CRISPRa and CRISPRi controls (lacking sgRNA) (One sample t-test, N=3). Synaptophysin was used as reference gene.

Next, we compared the modulatory effects of the two sgRNAs in both the CRISPRa and CRISPRi systems. A significant increase in *Cacna1h*-mRuby fluorescence activity was observed for the two single sgRNAs in the CRISPRa system (mean ± SEM= sgRNA1: 3.408 ± 0.258-fold increase, *p*=0.0026; sgRNA2: 2.925 ± 0.1571-fold increase, *p*=0.0012), and reached similar levels as observed for the combination of the two sgRNAs (**Fig 1D**, left panel; mean ± SEM= sgRNA1+2: 3.600 ± 0.5135-fold increase, *p*=0.0149). Also for the CRISPRi system, comparable modulatory effects were observed for sgRNA1 (mean ± SEM= 0.5250 ± 0.07773-fold decrease, *p*= 0.0088), sgRNA2 (0.5050 ± 0.04555-fold decrease, *p*=0.0017) and the combination of the two sgRNAs (0.5175 ± 0.1007-fold decrease, *p*= 0.0173; **Fig 1D**, middle panel). A control sgRNA targeting *Lac*Z did not have an effect on *Cacna1h* fluorescence intensity in the CRISPRa or CRISPRi system (**Fig 1D**, right panel), indicating that the modulatory effects observed for the two *Cacna1h*-sgRNAs were highly specific.

To confirm and precisely quantify the *Cacna1h*-specific CRISPRa and CRISPRi regulation, we next exchanged the mRuby reporter for a luciferase reporter gene (**Fig 1E**, left panel). As expected, transfection of the CRISPRa and CRISPRi components into NS20Y cells, resulted in an increase of luciferase activity for the CRISPRa system (**Fig 1E**, middle panel; sgRNA1: 5.438 ± 0.8392-fold increase, *p*=0.0339; sgRNA2: 4.789 ± 0.8021-fold increase, *p*=0.042; sgRNA1+2: 6.749 ± 0.4456-fold increase, *p*=0.006) and a decrease for the CRISPRi system (**Fig 1E**, right panel; sgRNA1: 0.2596 ± 0.08652-fold decrease, *p*=0.0134; sgRNA2: 0.2399 ± 0.03911-fold decrease, *p*=0.0026; sgRNA1+2: 0.1224 ± 0.01390-fold decrease, *p*=0.0003). Also here, no significant differences were observed between the two sgRNAs or the combination of the two sgRNAs.

We next examined the endogenous *Cacna1h* mRNA expression levels after manipulation with the CRISPRa and CRISPRi systems using quantitative real-time RT-PCR. Since both sgRNAs showed similar effects on the *Cacna1h* reporter constructs (**Fig 1D, E**) we decided to use the combination of the two sgRNAs for this experiment. Activation of the system in NS20Y cells, resulted in augmentation of endogenous *Cacna1h* mRNA expression levels, whereas inhibition significantly decreased the *Cacna1h* mRNA expression levels (**Fig 1F**, right panel; CRISPRa: 2.688 ± 0.3284 fold increase, *p*=0.0358; CRISPRi: 0.6898 ± 0.01801 fold decrease, *p*=0.0034). No changes in *Cacna1h* expression were observed for the CRISPRa and CRISPRi controls (no sgRNA; **Fig 1F**, left panel). Altogether, these results confirmed that the present *Cacna1h*-CRISPR modulatory toolbox successfully can activate or inhibit endogenous *Cacna1h* expression in NS20Y cells.

We then probed whether the modulatory effects observed in NS20Y cells could also be observed in primary cultured neurons. For this, we transduced primary mouse cortical neurons at day in vitro (DIV) 4 with all-in-one CRISPRa/i lentiviruses (**Fig 2A**) and measured endogenous *Cacna1h* mRNA expression levels at DIV15. Since the two sgRNAs displayed similar effects in NS20Y cells (**Fig 1D** and **E**), we decided to proceed with only one sgRNA (sgRNA2) to minimize unspecific effects in the neuronal cultures. Intriguingly, a strong activation (4.374 ± 0.3717-fold increase, *p*=0.0016) and inhibition (0.1233 ± 0.04931-fold decrease, *p*=0.0322) was observed after transduction with CRISPRa and CRISPRi lentiviruses, respectively (**Fig 2B**), indicating that sgRNA2 can efficiently modulate *Cacna1h* promoter activity in primary neurons.

**Figure 2.**
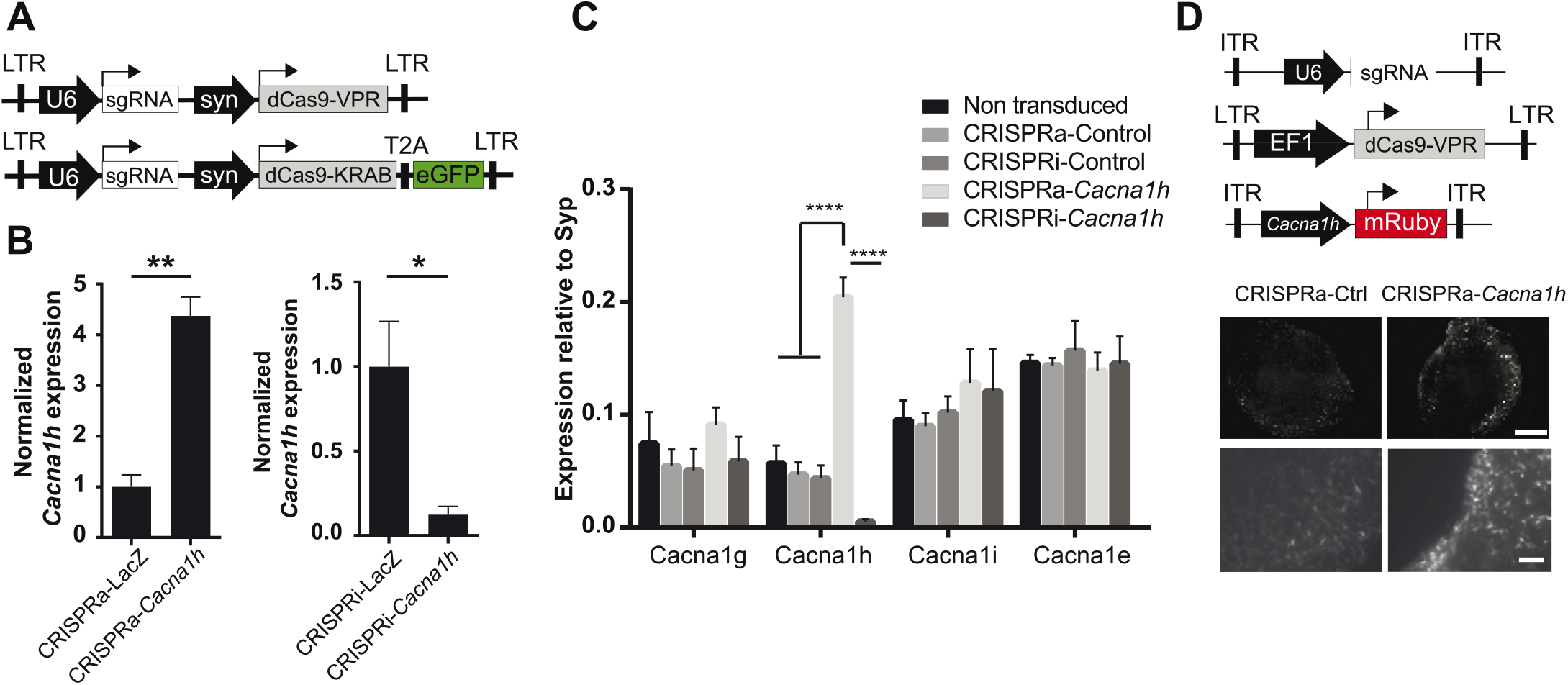
CRISPRa/i modulates the endogenous expression of *Cacna1h* in primary neurons. **(A)** Schematic representation of the all-in-one lentiviral constructs for CRISPRa (Lenti-U6-sgRNA-syn-dCas9-VPR) and CRISPRi (Lenti-U6-sgRNA-syn-dCas9-KRAB-EGFP). **(B)** mRNA expression of *Cacna1h* in cultured neurons after transduction with the all-in-one lentivirus targeting either *Cacna1h* or the control (*LacZ*). Synaptophysin was used as reference gene. Expression levels of *Cacna1h* relative to Synaptophysin in every treatment were normalized to the controls (CRISPRa-*LacZ* and CRISPRi-*LacZ*). Expression of *Cacna1h* was higher upon transduction with CRISPRa-*Cacnah1* in comparison to the control CRISPRa-*LacZ* (mean ± SEM: CRISPRa-*LacZ* 1.0 ± 0.2355; CRISPRa-*Cacnah1* 4.374 ± 0.3717, N=3, t-test, p=0.0016, **p≤0.01) and lower after transduction with CRISPRi-*Cacna1h* (mean ± SEM: CRISPRi-*LacZ* 1.0 ± 0.2673; CRISPRi-*Cacna1h* 0.1233 ± 0.04931, N=3, t test, p=0.0322, *p≤0.05). **(C)** mRNA expression of different calcium channels in neurons transduced with the all-in-one lentiviral constructs targeting *Cacna1h*. Expression relative to synaptophysin (N=3, Two way ANOVA, Tukey’s multiple comparison test, ****p≤0.0001). **(D)** Mouse organotypic hippocampal slices transduced with the CRISPRa lentivirus, the AAV-U6-sgRNA and the reporter AAV-*Cacna1h*-mRuby (Scale bar of 500 μM for upper and 100 µM for lower panel).

To prove unequivocally that the *Cacna1h*-CRISPRa/i toolbox is specific for *Cacna1h* modulation, we next examined the mRNA expression levels of other calcium channel family members (*Cacna1g, Cacna1i* and *Cacna1e*) after *Cacna1h*-CRISPRa/i targeting using quantitative real-time RT-PCR. No alterations were observed for any of the other calcium channels tested (**Fig 2C**), indicating that our newly developed *Cacna1h*-CRISPRa/i toolbox is highly specific for *Cacna1h* modulation.

In order to test if our system has the potential to be delivered in neuronal network structures *in vivo* we infected mouse organoptypic hippocampal slices with recombinant adeno-associated (rAAV) and lentiviruses encoding our *Cacna1h*-CRISPRa toolbox and a fluorescent reporter. After co-transduction with AAV-sgRNA, lenti-IF1adCas9-VPR (Savell et al., 2019) and the reporter AAV-*Cacna1h*-mRuby we observed a stronger signal of the red fluorescent protein when using the sgRNA targeting *Cacna1h* than in the control indicating an activation of the *Cacna1h* promoter *in vivo* (**Fig 2D**).

## 4 Discussion

Detailed analyses of the dynamic contribution of individual ion channels in the context of neuronal network function remains challenging, particularly in the context of mutational and transcriptional ‘channelopathies’, in which for example the fine-tuned regulation of mRNA expression levels of affected genes, including *Cacna1h* plays a major role. Traditionally, the functional characterization of individual ion channels in terms of gain- and loss-of-function approaches has been addressed by the exogenous overexpression of genetically encoded proteins and by knock-down systems via RNA interference (RNAi) / genetic gene ablation (knock-out), respectively. Although very valuable, the applicability of these techniques has limitations (Kampmann, 2018).

Here, we present a modular method using a CRISPRa/i system that allows to manipulate the endogenous expression of *Cacna1h in-vitro* and *in-vivo*. This CRISPRa/i system applies specific sgRNAs directed to the *Cacna1h* promoter and a dCas9 fused to the transcriptional activator VPR or the transcriptional inhibitor KRAB (Gilbert et al., 2013; Chavez et al., 2015) to induce changes in the endogenous expression of the *Cacna1h* gene. We provide evidence that both the CRISPRa and CRISPRi systems can specifically manipulate the endogenous expression of *Cacna1h* in dividing cells as well as in primary neuronal cultures. In addition, we demonstrate the possibility to activate the *Cacna1h* promoter in mouse organotypic hippocampal slices. Interestingly, the fold-change induction by CRISPRa occurred within a range also observed for Ca_V_3.2 channelopathies (Becker et al., 2008), making our *Cacna1h*-CRISPR system highly suitable for analyzing this particular and also other channelopathies at the functional level.

Increasing the gene expression levels of *Cacna1h* in a high percentage of cultured cells, which is required for many downstream analyses, or *in-vivo* requires the application of viral transduction systems. Here, our *Cacna1h*-CRISPRa toolbox has a clear advantage over conventional virally-mediated overexpression approaches, where the limiting packaging capacity of the commonly used and easily applicable recombinant adeno-associated- and lenti-viruses, excludes the possibility to overexpress large proteins like *Cacna1h*. On the other hand, recombinant viruses with a higher packaging capacity, like Adeno- and Herpes Simplex virus, cannot be easily generated in the lab and are expensive to purchase. Since augmentation of proteins using the CRISPRa system is independent of their transcript size, even the large *Cacna1h* gene (mRNA transcript length up to 8.2 Kb) can be easily augmented using rAAV and lentiviruses, which can be produced in most molecular biology labs. Another advantage of the *Cacna1h*-CRISPRa system over conventional gain-of-function approaches is the endogenous modulation of the *Cacna1h* genomic locus, providing the opportunity to study the effects of abundant alternative splicing variants of *Cacna1h*. For *Cacna1h*, at least 12-14 alternative splicing sites have been described, resulting in the possibility to generate more than 4000 alternative *Cacna1h* transcripts (Zhong et al., 2006). Such a complexity cannot be achieved by conventional overexpression of only one variant of *Cacna1h*. Also in terms of loss-of-function approaches, our *Cacna1h*-CRISPRi has clear advantages over conventional shRNA/siRNA approaches. A siRNA approach for *Cacna1h* in mouse embryonic stem cells reduced *Cacna1h* expression levels to approximately 40% but also caused a non-specific decrease in *Cacna1g* mRNA expression levels (Rodríguez-Gómez et al., 2012). The fact that we did not observe any alterations in the mRNA expression levels of the calcium channel family members *Cacna1g, Cacn1i* and *Cacna1e* clearly indicates that our newly developed *Cacna1h*-CRISPRa/i toolbox is highly specific for *Cacna1h* modulation. Importantly, since the catalytic inactive Cas9 (dCas9) used in our system does not cleave the DNA, the possible off-target effects are more likely to be less deleterious than using the conventional Cas9 to induce a genetic knock-out (Colasante et al., 2020). Compared to genetic knock-out approaches, the *Cacna1h*-CRISPRa/i toolbox, which is based on viral-transduction, allows to decrease *Cacna1h* expression in selected neuron types, by putting dCas expression under the control of cell type specific promoters, in localized brain regions and under temporal control, without time consuming breeding of animals.

The *Cacna1h*-CRISPR toolbox we describe here is made for constitutive activation or inhibition of the *Cacna1h* transcripts. Nevertheless, the system could easily be adapted for induced gene expression control by incorporating Doxycycline-controlled Tet-On and Tet-Off gene expression systems (Colasante et al., 2019; Zhang et al., 2019). In this way, *Cacna1h* expression can be activated or inhibited at a precise temporal resolution. The possibility of combining the C*acna1h-*CRISPR toolbox with a Tet-On/Off inducible system might open new roads for investigating the role of *Cacna1h* in several channelopathies in more detail. For example, is a transient augmentation of *Cacna1h* alone sufficient to elicit an epileptic phenotype? A transient increase of *Cacna1h* has been observed early after pilocarpine-induced SE and knockout mice for *Cacna1h* show substantially attenuated development of spontaneous chronic seizure activity (Becker et al., 2008). From these data, it was concluded that a transient *Cacna1h* augmentation, representing a ‘transcriptional channelopathy’, is critical for the process of epileptogenesis. In this regard, an intriguing analysis option is given by using a Tet-On C*acna1h-*CRISPR approach in order to test whether a transient increase of *Cacna1h* alone results in spontaneous seizure activity or whether additional factors are mandatory (**Fig 3A**). Additionally, the Tet-inducible C*acna1h-*CRISPR system could also be used to reveal the specific time window for *Cacna1h* upregulation to be important during the process of epileptogenesis. For this, the C*acna1h-*CRISPRi can be incorporated with the Tet-on system, providing the possibility to temporally inhibit *Cacna1h* expression at specific time-intervals early after pilocarpine-induced SE and to measure the phenotypic outcome (**Fig 3B**).

**Figure 3.**
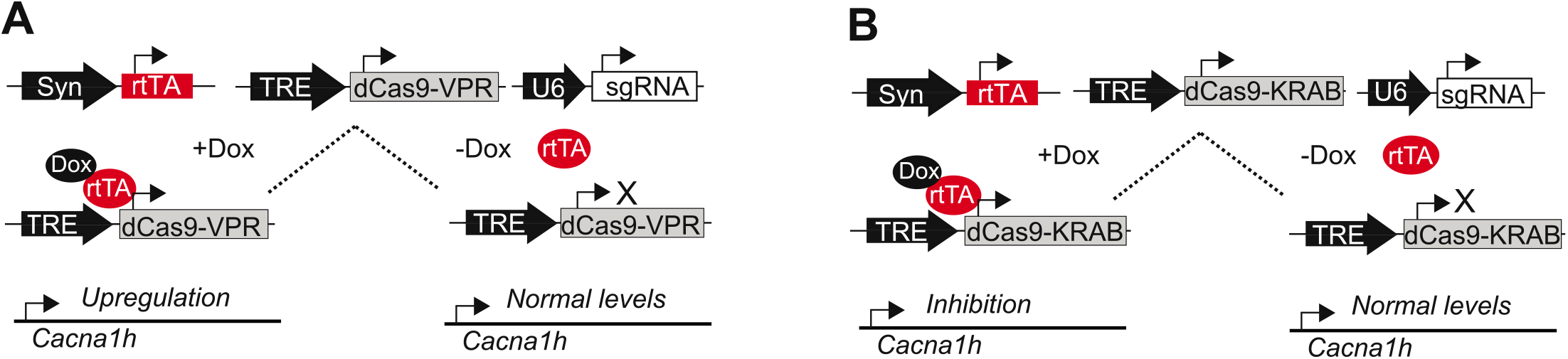
Future perspectives to study *Cacna1h*-channelopathies. **(A)** Schematic representation of a TET-ON system for the conditional expression of CRISPRa (TRE-dCas9-VPR). Only in the presence of doxycycline (Dox) the reverse tetracycline-controlled transactivator (rtTA) binds to the tetracycline response element (TRE) inducing the expression of *Cacna1h*-CRISPRa resulting in augmented endogenous *Cacna1h* expression. **(B)** Schematic representation of a TET-ON system for the conditional expression of CRISPRi (TRE-dCas9-KRAB). Only in the presence of Dox, endogenous *Cacna1h* expression is inhibited.

Also in the context of mutational/genetic ion channelopathies, the potential of inducible *Cacna1h*-CRISPR approaches are immense. For example, in the Genetic Absence Epilepsy Rat from Strasburg (GAERS) a mutation in C*acna1h* has been described, which augments the expression of Ca_v_3.2 at the cell surface and increases calcium influx (Proft et al., 2017). Here, the *Cacna1h-*CRISPRa could be used as a gain-of-function approach to mimic the enhanced expression seen in GEARS. In addition, a transient inhibition of *Cacna1h* via the Tet-on inducible *cacna1h-*CRISPRi system in the GAERS rats at different stages during development could also reveal potentially interesting time windows for *Cacna1h-*associated channelopathies **(Fig 3B)**.

Taken together, our here described *Cacna1h*-CRISPRa/i modular approach could thus be used to model transient gain-of-function or loss-of-function effects in the *Cacna1h* gene and to study Ca_V_3.2 channelopathies in more detail *in-vivo*.

## 5 Ethical Publication Statement

We confirm that we have read the Journal’s position on issues involved in ethical publication and affirm that this report is consistent with those guidelines.

## 7 Author Contributions

DT, AJB and KMJvL conceived and planned the study. DT and ART carried out molecular and cellular experiments. HS and SS contributed to study design and analysis. DT, AJB, SS and KMJvL wrote the manuscript.

## 9 Acknowledgments

We thank Sabine Optiz for excellent technical assistance.

## 10 Data Availability Statement

The raw data supporting the conclusions of this article will be made available by the authors, without undue reservation.

